# Circulating miR-4532 is associated with loss of ambulation in dysferlinopathy

**DOI:** 10.64898/2026.01.05.697793

**Authors:** Tajveer S. Grewal, Zsuzsanna Hollander, Darlene LY Dai, Virginia Chen, Hillarie P. Windish, Douglas E. Albrecht, Elaine Lee Paoliello, Laura Rufibach, Bradley Williams, Plavi Mittal, Sara Assadian, Janet E. Wilson-McManus, Bruce McManus, Raymond Ng, Scott Tebbutt, Pascal Bernatchez, Amrit Singh

## Abstract

**Background:** Limb-girdle muscular dystrophies (LGMDs) are inherited myopathies characterized mainly by progressive weakness of the proximal muscles of the shoulder and pelvic girdle areas, leading to functional decline and eventual loss of independent ambulation. Dysferlinopathy (LGMD2B) is an autosomal recessive LGMD subtype, is caused by mutations in the *DYSF* gene that lead to lack of dysferlin which results in muscle death and chronic muscle fiber degeneration. Preservation of ambulation is a key clinical milestone, as loss of independent gait markedly reduces quality of life and complicates care management. Although patients often perceive functional decline before their initial clinical presentation, current clinical assessments typically detect disease progression after substantial muscle damage has occurred.

**Methods:** In this multigroup case-control study, we profiled plasma miRNAs from 49 genetically confirmed dysferlinopathy patients (24 ambulatory, 25 non-ambulatory) and 25 age- and sex-matched healthy controls. Total RNA was extracted from blood samples and hybridized to Affymetrix GeneChip miRNA 3.1 arrays. After quality control and filtering, differential expression analysis was performed using linear models for microarrays, adjusting for age and sex, with a false discovery rate cutoff of 10%.

**Results:** 14 miRNAs were significantly altered between dysferlinopathy patients and controls. Notably, miR-4532 was upregulated in ambulatory patients relative to controls, whereas it was downregulated in non-ambulatory patients compared with ambulatory patients, although expression levels remained higher than in controls. Levels of miR-4532 were positively associated with circulating monocyte levels in ambulatory patients only.

**Conclusion:** These results suggest that miR-4532 may be a circulating marker associated with ambulatory status in dysferlinopathy. Its known involvement in inflammatory signaling and muscle regeneration pathways underscores its potential as an early indicator for disease activity.

## Introduction

Muscular dystrophies (MDs) are a heterogenous group of inherited genetic disorders characterized by progressive muscle weakness and degeneration (1). A large number of distinct MDs have been identified, each caused by mutations in genes encoding muscle proteins (e.g., dystrophin in Duchenne MD) (1). MDs share common features of skeletal muscle fiber wasting and weakness, but differ in age of onset, severity, and the specific muscle group affected (1,2). Limb-girdle muscular dystrophies (LGMDs) are 1 major subclass with over 30 different genetically identified subtypes, defined by early involvement of shoulder and pelvic girdle muscles (proximal legs, too). LGMD is described as a “progressive weakness and atrophy of the hip, shoulder, and proximal extremity muscles”, often beginning in adolescence or early adulthood (2). Cardiac or respiratory involvement is variable depending on subtype.

Dysferlinopathy (LGMD2B) is an autosomal recessive form of LGMD caused by mutations in the *DYSF* gene which results in dysferlin deficiency (2). Dysferlin encodes a ∼237 kDa transmembrane protein in the sarcolemma, rich in Ca^2+^-binding C2 domains, that regulates Ca^2+^-dependent vesicle fusion (3,4). Loss of dysferlin impairs the normal resealing of muscle cell membranes after injury, and also causes aberrant Ca^2+^ and lipid handling (3–6). Dysferlinopathy encompasses multiple clinical phenotypes, including both proximal limb-girdle (LGMD2B/R2) and distal Miyoshi Myopathy type 1 (MMD1) presentations; which have been shown to be the same disease (7). Onset of dysferlinopathy usually occurs in late adolescence or early adulthood, initially as subtle weakness (for example, loss of ankle reflexes and difficulty standing on tiptoes) and then slowly progresses to involve more proximal and distal limb muscles (3).

The manifestations of dysferlinopathy are clinically significant because they cause debilitating proximal muscle weakness and functional decline. Patients typically have symmetric weakness of pelvic and shoulder girdle muscles, resulting in difficulty climbing stairs, rising from chairs, or lifting objects (2). Over time, muscle degeneration leads to gait instability and progressive loss of function (8). Thus, preserving ambulation is a key treatment goal in the care management for dysferlinopathy.

Because progression of dysferlinopathy is slow and heterogenous, there is great interest in identifying non-invasive biomarkers that can detect disease activity before major functional loss. Circulating blood biomarkers including proteins, metabolites, DNA, and microRNAs have been widely explored for MDs, since they can be measured longitudinally to monitor disease progression or therapeutic response (9). Recent studies have shown that specific plasma miRNAs are differentially expressed in MD patients (10). In a survey of LGMD patients, 13 circulating miRNAs distinguished patients from controls (10). Such findings suggest circulating miRNAs might serve as sensitive indicators of muscle pathology. Notably, a larger study that included patients across multiple disease stages demonstrated distinct miRNA expression patterns across several muscular dystrophies, supporting the robustness and generalizability of miRNA dysregulation in muscle disease (11). However, few studies have correlated specific miRNA changes with clinical status in patients with LGMDs.

In this study we aimed to identify circulating miRNAs whose levels differ between healthy controls and dysferlinopathy patients at different stages of the disease. By profiling these miRNAs in blood samples from dysferlinopathy patients with varying ambulatory status, we aim to identify early molecular indicators of disease progression. The long-term goal is to use identified biomarkers to enable earlier intervention while patients are still ambulatory, thereby helping to preserve mobility and function in dysferlinopathy.

## Materials and Methods

### Patient enrollment

Fifty genetically confirmed dysferlinopathy participants were prospectively enrolled between April 2014 and January 2015 throughout the United States of America (H13-023848). The study examined samples from 2 groups at the extreme ends of the phenotypic spectrum: 25 dysferlinopathy participants with advanced disease (non-ambulatory) and 25 dysferlinopathy participants who were highly ambulatory and therefore considered to be in an early disease stage. Ambulation status of dysferlinopathy patients was assessed using an online quiz in which patients responded with their current ability to walk and run, noting whether they were assisted and unassisted. A control group of 25 healthy participants were enrolled in Vancouver, Canada, chosen to reflect similar marginal age and sex distributions as the dysferlinopathy participants.

### Blood collection

Blood samples were collected from each of the 75 participants using PAXgene tubes. PAXgene blood was kept on cold gel packs after collection and during shipment to the St. Paul’s Hospital clinical laboratory in Vancouver, Canada. Once samples were received in Vancouver, they were frozen at -80°C. The processing of control participants’ blood was analogous to the dysferlinopathy participant blood. Complete blood counts and differentials were obtained for blood samples when possible.

### Molecular profiling

Microarray analysis on PAXgene samples was performed at the TSRI DNA Array Core Facility, Scripps Research Institute (La Jolla, CA). Total RNA was extracted on the QIAcube (Qiagen Inc) from PAXgene blood tubes using the PAXgene Blood miRNA kit from PreAnalytix (Cat. #763134) according to the manufacturer’s instructions. The miRNA analysis was performed using the total RNA extracted for the PAXgene tubes. Samples were hybridized overnight to Affymetrix GeneChip miRNA 3.1 arrays.

### Statistical analysis

All analyses were performed using the statistical computing software R (v4.4.1). ANOVA analysis was used to determine which factors (group, RNA extraction batch, sex, and age) were associated with the miRNA expression profiles represented by using Principal component analysis. Linear Models for Microarrays (LIMMA, v3.60.6) was used to perform differential miRNA analysis, adjusted for age and sex. Three pairwise comparisons were made between groups: controls versus ambulatory, controls versus non-ambulatory and ambulatory versus non-ambulatory. P-value correction for multiple hypothesis testing was performed using the Benjamini-Hochberg false discovery rate (BH-FDR) and BH-FDR threshold of 10% was used to identify differentially expressed miRNAs. Gene targets of select miRNAs were identified using miRNet 2.0 and used for gene ontology analysis using Enrichr (enrichR, R-package v3.2) (12,13).

## Results

### Patient characteristics

The cohort consisted of 75 subjects (Table 1). Of these, 50 had a confirmed diagnosis of a dysferlinopathy based on the presence of mutations in the *DYSF* gene and had a clinical presentation of either LGMD2B or MMD1 (collectively called dysferlinopathy). Twenty-five of the patients were ambulatory and the other 25 had more advanced disease (wheelchair bound; non-ambulatory). On average, non-ambulatory patients were older (∼49y) than ambulatory patients (∼33y). The remaining 25 discovery cohort subjects consisted of healthy controls (average age of 37y).

**Table 1.**
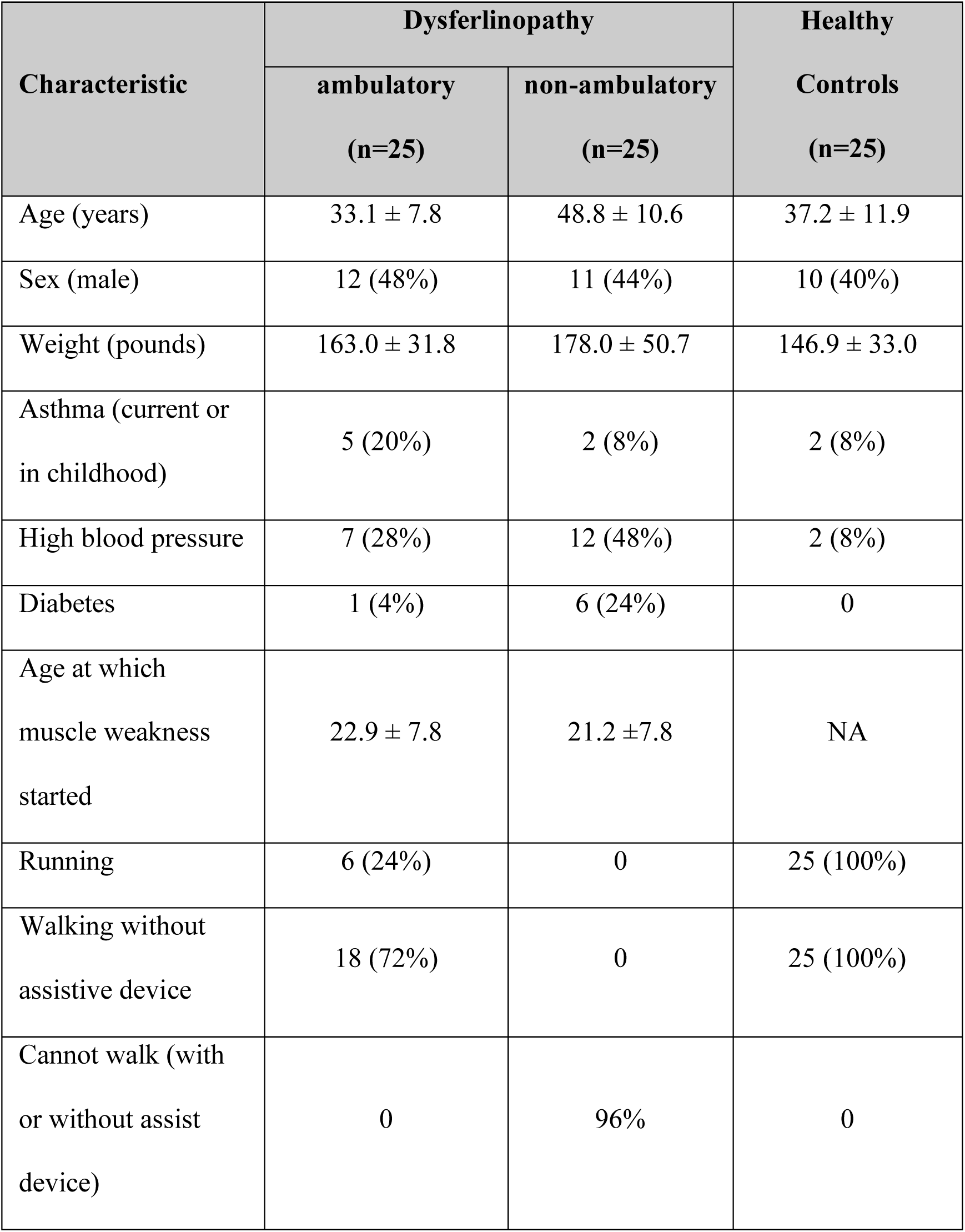
Patient characteristics.

| Characteristic | Dysferlinopathy |  | Healthy Controls<br>(n=25) |
| --- | --- | --- | --- |
|  | ambulatory<br>(n=25) | non-ambulatory<br>(n=25) |  |
| Age (years) | 33.1 ± 7.8 | 48.8 ± 10.6 | 37.2 ± 11.9 |
| Sex (male) | 12 (48%) | 11 (44%) | 10 (40%) |
| Weight (pounds) | 163.0 ± 31.8 | 178.0 ± 50.7 | 146.9 ± 33.0 |
| Asthma (current or in childhood) | 5 (20%) | 2 (8%) | 2 (8%) |
| High blood pressure | 7 (28%) | 12 (48%) | 2 (8%) |
| Diabetes | 1 (4%) | 6 (24%) | 0 |
| Age at which muscle weakness started | 22.9 ± 7.8 | 21.2 ± 7.8 | NA |
| Running | 6 (24%) | 0 | 25 (100%) |
| Walking without assistive device | 18 (72%) | 0 | 25 (100%) |
| Cannot walk (with or without assist device) | 0 | 96% | 0 |

### Loss of ambulation results in more differentially expressed miRNAs in dysferlinopathy patients compared to controls

One sample (ambulatory subject) had a poor-quality miRNA array and was excluded from further analysis. Of the 6,658 miRNAs profiled, miRNAs that were either unannotated or had an average expression across all 74 samples of less than 3 units were removed. This resulted in 666 miRNAs (45 “mir” or pre-miRNAs, 439 “miR” or mature miRNAs, 11 “let” miRNAs of the let family, and 171 small nucleolar RNA [snoRNAs]). Principal component analysis was applied to the resulting miRNA dataset, with the first 5 principal components (PCs) capturing ∼60% of the variation. **Figure 1A**, depicts the significance of the association between each PC and factors such as disease group, RNA extraction batch, sex and age. PC2 and PC4 were strongly associated with disease group and sex, respectively, whereas PC1 and PC2 were modestly associated with age, suggesting an influence of sex and age on the miRNA expression profiles. No significant association with RNA extraction batch was observed. A linear model was used to regress the expression of each miRNA onto the groups (controls, ambulatory and non-ambulatory), age and sex. The strongest effect was observed when comparing controls with non-ambulatory patients (green line in **Figure 1B**), whereas the weakest differential signal was observed when comparing between ambulatory and non-ambulatory patients (blue line in **Figure 1B**). At a BH-FDR of 10%, miR-4532 was significantly up-regulated in ambulatory patients as compared to controls. At the same BH-FDR threshold, miR-4532 and ACA40 (a snRNA) were up-regulated in non-ambulatory patients compared to controls, whereas 9 miRNAs were down-regulated in non-ambulatory patients compared to controls (**Figure 1C**). No significant miRNAs were identified between ambulatory and non-ambulatory patients at a BH-FDR threshold of 10%.

**Figure 1.**
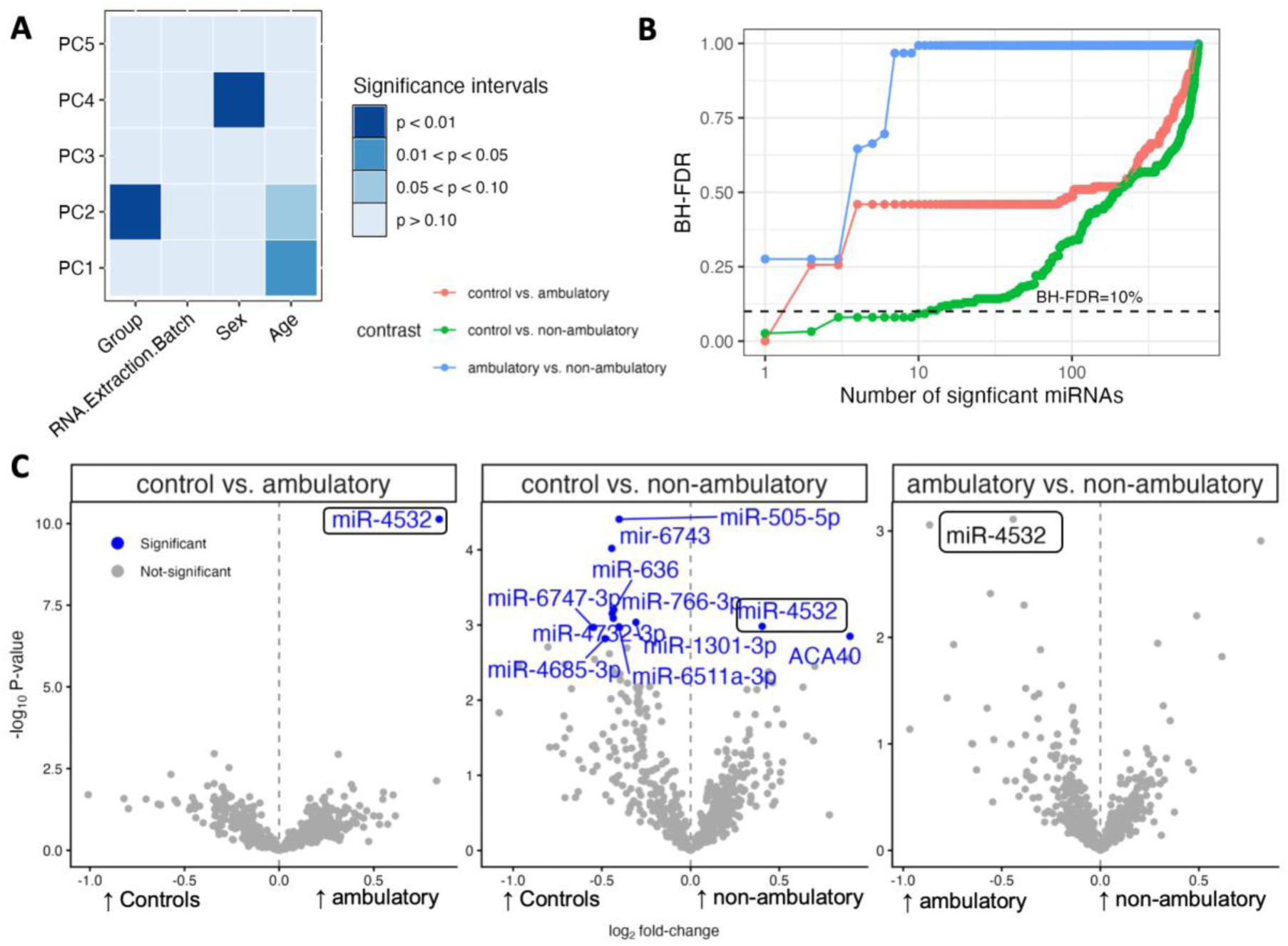
Differential miRNA expression across dysferlinopathy disease states. **(A)** Association of principal components (PC1–PC5) with clinical and demographic variables, showing the contribution of group status (control, ambulatory, non-ambulatory), sex, and age to global miRNA expression variability. **(B)** Cumulative number of significantly differentially expressed miRNAs identified across Benjamini–Hochberg false discovery rate (BH-FDR) thresholds for each pairwise comparison. The dashed line indicates the BH-FDR=10% cutoff. **(C)** Volcano plots depicting differential miRNA expression for control vs. ambulatory, control vs. non-ambulatory, and ambulatory vs. non-ambulatory dysferlinopathy patients. Highlighted miRNAs in blue meet the BH-FDR=10% threshold. All analyses were adjusted for age and sex.

### Association of miR-4532 with loss of ambulation in dysferlinopathy patients

Limiting the differential expression analysis for the ambulatory and non-ambulatory comparison to the 11 significant miRNAs (found when comparing controls vs. ambulatory patients), resulted in miR-4532 as the only statistically differentially expressed miRNA (*p*=0.00078, BH-FDR_new_=0.9%, BH-FDR_old_=27.6%). **Figure 2A** shows that miR-4532 is up-regulated in dysferlinopathy regardless of ambulation status, however, the levels of miR-4523 were significantly reduced in non-ambulatory patients compared to ambulatory patients. Given that these miRNA expression profiles can be attributed to immune cells, we tested the association between miR-4532 with various complete blood counts and differentials. **Figure 2B** shows a positive association with monocytes in ambulatory patients and a negative correlation with neutrophils and various red blood counts in non-ambulatory patients, suggesting that miR-4532 is likely expressed by monocytes in ambulatory patients. **Figure 2C** depicts a lack of association between monocytes and miR-4532 in controls, a significant positive association in ambulatory patients (*p*=0.038) and negative association in non-ambulatory patients. 148 gene-targets were identified for miR-4532 using miRNet 2.0 and 2 significant gene ontology (GO) molecular functions were identified (BH-FDR=10%): 4-way junction DNA binding and phosphatidylionositol-3,5-bisphosphate binding (**Figure 2D**).

**Figure 2.**
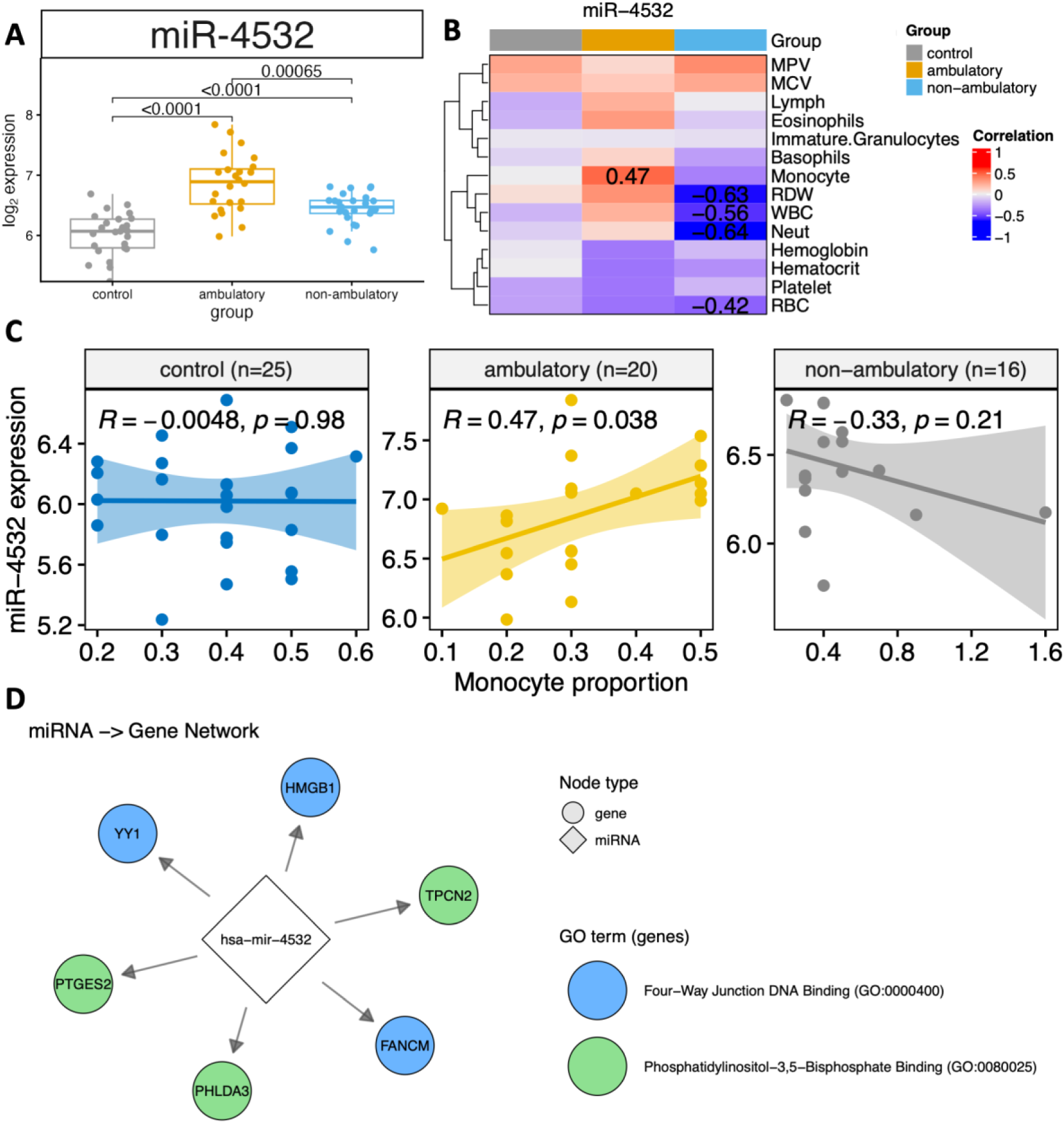
MiR-4532 discriminates between controls and dysferlinopathy patients. A. Boxplot of miR-4532 expression of in controls and dysferlinopathy patients with and without loss of ambulation. B. Heatmap of Pearson correlations for complete cell counts and differentials with miR-4532 expression in each group. Only values greater than |0.40| are displayed. C. Scatter plot between miR-4532 expression and monocyte proportions per group. The number of samples with available monocytes counts are depicted by n for each group. D. Network of miR-4532 and its gene-targets based on Gene Ontology analysis.

## Discussion

In the present study we demonstrate the up-regulation of circulatory levels of miR-4532 in dysferlinopathy patients as compared to healthy controls. The levels of miR-4532 were associated with monocyte levels in ambulatory patients only which may indicate a cell-specific role for miR-4532 that changes over the course of the disease.

miR-4532 has not been previously linked to MD but has been identified in settings of inflammatory and regeneration. In immune-related studies, miR-4532 emerges repeatedly. For example, urinary exosome profiling showed a 3-miRNA signature including miR-4532 that distinguished kidney transplants with acute rejection (immune-mediated injury) from stable graft recipients (14). Mechanistically, macrophage exosomes rich in miR-4532 can provoke inflammation: 1 study found that miR-4532 in macrophage-derived exosomes binds to the 3’-untranslated region (UTR) of specificity protein 1 (SP1), triggering SP1 degradation and downstream NF-κB (p65) activation (15). This SP1/NF-κB signaling drives endothelial inflammation and injury. Thus, high exosomal miR-4532 promotes pro-inflammatory signaling via NF-κB (16).

In dysferlinopathy, chronic muscle fiber necrosis elicits macrophage infiltration; by analogy, miR-4532 might modulate NF-κB – driven immune responses. However, no direct data exists on miR-4532 in dystrophic muscle or muscle macrophages. The observation that miR-4532 is upregulated in dysferlinopathy patients compared to healthy controls yet downregulated in non-ambulatory versus ambulatory patients is demonstrated in **Figure 2A**. Compared to miR-4532 in other diseases, this presents a seemingly paradoxical pattern that warrants careful interpretation (16). The expression profile suggests that miR-4532 levels peak during the early, ambulatory stages of the disease and then decline as the disease progresses to the non-ambulatory stage. Rather than indicating a worsening pathogenic state, the decrease in miR-4532 levels in non-ambulatory patients likely reflects reduced muscle mass and potentially a less active inflammatory state.

Based on the gene ontology analysis revealed in **Figure 2D**, miR-4532 is significantly associated with 4-way junction DNA binding. 4-way junction DNA binding is a common property of architectural proteins involved in DNA repair, recombination, and chromatin remodeling. The network analysis identifies three key target genes of miR-4532 that participate in 4-way junction DNA binding: Ying Yang 1 (*YY1)*, High Mobility Group Box 1 (*HMGB1)*, and Fanconi Anemia Complementation Group M *(FANCM)*. *YY1* is a ubiquitously expressed transcription factor that regulates both transcriptional activation and repression (17). *YY1* forms protein complexes with the BAF chromatin remodeling complex and targets both promoters and super-enhancers to activate transcription of pluripotency genes (17). Its binding to 4-way DNA junctions enables it to recognize structural distortions in DNA and manipulate DNA structure during transcriptional regulation (17). *HMGB1* is a nonhistone chromatin-associated protein that binds to 4-way DNA junctions in their open conformation (18). Further, *HMGB1* selectively binds to structural RNAs, particularly H/ACA box snoRNAs and scaRNAs, and regulates ribosomal RNA processing (18). The ability of *HMGB1* to bind 4-way junctions makes it crucial for chromatin organization and transcriptional regulation. *FANCM* represents the most sophisticated target, functioning as both a DNA translocase and a sensor for the Fanconi Anemia pathway (19). *FANCM* employs a dual recognition mechanism using distinct DNA-binding domains to identify and remodel branched DNA structures at damaged sites (19). Its N-terminal translocase domain specifically recognizes branched DNA through Hel2i and “mincer” domains, while coupling this recognition to Faconi Anemia pathway activation through recruitment of the Faconi Anemia core complex (19).

The second major functional category associated with miR-4532 involves phosphatidylinositol 3,5-biphosphate binding, targeting genes Two Pore Segment Channel 2 (*TPCN2*), Prostaglandin E Synthetase 2 (*PTGES2)*, and Pleckstrin Homology-Like Domain Family A, Member 3 (*PHLDA3)*. This represents a novel connection between miR-4532 and lipid signaling pathways that regulate membrane dynamics and cellular metabolism. *TPCN2* is an intracellular Ca^2+^-release channel located to endolysosomal membranes (20). *TPCN2* functions as a sensor activated by both nicotinic acid adenine dinucleotide phosphate (NAADP) and phosphatidylinositol 3,5-bisphosphate (21). The channel is highly Ca^2+^-selective and its gating is regulated by luminal pH and Mg^2+^, making it crucial for endolysosomal Ca^2+^ signaling (21). *TPCN2* has been implicated in diverse cellular processes including virus entry, angiogenic signaling, and pigmentation regulation (21). *PTGES2* catalyzes the conversion of prostaglandin H2 to prostaglandin E2, a key inflammatory mediator (22,23). The enzyme is membrane-associated and glutathione-dependent, with a critical cysteine residue essential for enzymatic activity (22,23). Aberrant PTGES2 expression has been implicated in various cancers, with overexpression correlating with higher tumor grade and poorer prognosis (22,23). PHLDA3 is a p53-regulated tumor suppressor protein that contains a pleckstrin homology domain (24). PHLDA3 binds to various phosphatidylinositol phosphates through this domain and localizes to cell membranes (24). The protein functions as a domain-negative repressor of AKT signaling by competing with AKT for PIP3 binding on the plasma membrane, thereby inhibiting AKT activation and downstream pro-survival signaling (24). PHLDA3 is frequently downregulated in various cancers through various mechanisms including loss of heterozygosity and promoter methylation (24).

The connection between miR-4532 and both DNA repair machinery (through 4-way junction binding targets) and lipid signaling pathways (through phosphatidylinositol binding targets) suggests that this microRNA may serve as a critical regulatory hub linking genomic stability maintenance with membrane homeostasis and inflammatory responses. In the context of dysferlinopathy, where both membrane repair defects and chronic inflammation are prominent features, miR-4532’s regulation of these pathways may contribute significantly to disease progression.

The identification of miR-4532 as a differentially expressed miRNA in dysferlinopathy has several potential clinical implications. One of the main possible uses of miR-4532 levels would be for disease monitoring. Longitudinal monitoring of miR-4532 levels in patients could provide valuable insights into disease activity and help guide clinical management decisions. Changes in miR-4532 expression might precede clinical deterioration, allowing for earlier intervention. Unlike creatine kinase, which is a general marker of muscle damage, miR-4532 may provide more specific information about the inflammatory and regenerative status of the muscle of patients with dysferlinopathy. Similar to the decrease in creatine kinase levels with advancing disease, the reduction in miR-4532 observed with disease progression is likely attributable to progressive loss of muscle mass.

A clear limitation of the present study is the lack of cellular specificity of miR-4532. Although our results indicate that it is potentially expressed by macrophages, future studies that isolate macrophages at various stages of dysferlinopathy are needed to validate that miR-4532 as a macrophage-specific biomarker. Another limitation is the significantly older age of the non-ambulatory patients compared to ambulatory patients. Although we adjusted for age in the linear models, it is unknown whether miR-4532 typically increases or decreases with age.

Overall, the differential expression of miR-4532 offers valuable insights into the complex pathophysiology of dysferlinopathy. The established connection between miR-4532, macrophage function, and NF-kB signaling provides a compelling framework for understanding how this miRNA might contribute to the inflammatory processes that characterize dysferlinopathy. Further research is needed to further elucidate the specific targets and functions of miR-4532 in muscle and immune cells, as well as its potential utility as a biomarker and therapeutic target in dysferlinopathy. By integrating our understanding of miR-4532 with the broader context of dysferlinopathy pathogenesis, we can develop a more comprehensive approached to monitoring and treating this debilitating condition, ultimately improving outcomes for patients with dysferlinopathy.

## Acknowledgments

We thank the patients who gave consent for the study to use their samples and data. The Jain Foundation provided not only funding, but also helped in the design, implementation, analysis, review of the study, and patient recruitment. We would like to thank Dr. Steve Head and team from the Scripps Research Institute for assistance with the molecular analyses. The introduction and discussion sections were edited and paraphrased with the assistance of ChatGPTv4o (OpenAI). The authors verified the content prior to incorporation into the main text.

